# RubyACRs, non-algal anion channelrhodopsins with highly red-shifted absorption

**DOI:** 10.1101/2020.04.15.043158

**Authors:** Elena G. Govorunova, Oleg A. Sineshchekov, Hai Li, Yumei Wang, Leonid S. Brown, John L. Spudich

**Author notes:** Address correspondence to: John L. Spudich. **Author Contributions** E.G.G., O.A.S., L.S.B. and J.L.S. conceptualized the study. E.G.G., O.A.S., H.L. and Y.W. carried out experiments. All authors discussed and analyzed the results and contributed to writing of the manuscript.

## Abstract

Channelrhodopsins are light-gated ion channels widely used to control neuronal firing with light (optogenetics). We report two previously unknown families of anion channelrhodopsins (ACRs), one from the heterotrophic protists labyrinthulomycetes and the other from haptophyte algae. Four closely related labyrinthulomycete ACRs, named RubyACRs here, exhibit a unique retinal binding pocket that creates spectral sensitivities with maxima at 590-610 nm, the most red-shifted channelrhodopsins known, long-sought for optogenetics, and more broadly the most red-shifted microbial rhodopsins so far reported. We identified three spectral tuning residues critical for the red-shifted absorption. Photocurrents recorded from the RubyACR from *Aurantiochytrium limacinum* (designated *Al*ACR1) under single-turnover excitation exhibited biphasic decay, the rate of which was only weakly voltage-dependent, in contrast to that in previously characterized cryptophyte ACRs, indicating differences in channel gating mechanisms between the two ACR families. Moreover, in *A. limacinum* we identified three ACRs with absorption maxima at 485, 545, and 590 nm, indicating color-sensitive photosensing with blue, green and red spectral variation of ACRs within individual species of the labyrinthulomycete family. We also report energy transfer from a cytoplasmic fluorescent protein domain to the retinal chromophore bound within RubyACRs, not seen in similar constructs in other channelrhodopsins.

**Significance Statement:** Our identification and characterization of two ACR families, one from non-photosynthetic microorganisms, shows that light-gated anion conductance is more widely spread among eukaryotic lineages than previously thought. The uniquely far red-shifted absorption spectra of the subset we designate RubyACRs provide the long-sought inhibitory optogenetic tools producing large passive currents activated by long-wavelength light, enabling deep tissue penetration. Previously only low-efficiency ion-pumping rhodopsins were available for neural inhibition by the orange-red region of the spectrum. The unusual amino acid composition of the retinal-binding pocket in RubyACRs expands our understanding of color tuning in retinylidene proteins. Finally, energy transfer from the fluorescent protein used as a tag on RubyACRs opens a potential new dimension in molecular engineering of optogenetic tools.

## Introduction

Channelrhodopsins are light-gated ion channels, first found in chlorophyte (green) flagellate algae as phototaxis receptors that depolarize the cell membrane by cation conduction (1-3). The genomes of cryptophyte algae encode a related family of channelrhodopsins that are, however, strictly anion-selective (4). Both channelrhodopsin families, known as cation and anion channelrhodopsins (CCRs and ACRs, respectively), are widely used to, respectively, activate (5) and inhibit (6) neurons with light (optogenetics).

Recently a distinct family of ACRs was identified in environmental DNA samples of unknown organismal origin collected by the Tara Oceans project (7). This finding raised a possibility that channelrhodopsin genes were not limited to algae and were more widespread among eukaryotic protists. Here we report two distinct ACR families from labyrinthulomycetes and haptophytes. Labyrinthulomycetes are a class of aquatic saprotrophic microbes included in a larger group called stramenopiles (8). Labyrinthulomycete ancestors are believed to have never been photosynthetic (9), unlike some other heterotrophic eukaryotic lineages that evolved by plastid loss (e.g. Apicomplexa). Haptophytes are flagellate algae distantly related to cryptophytes (10), but nevertheless segregated to the separate Haptista supergroup (11).

Long-wavelength light better penetrates biological tissue, and therefore actuator molecules with red-shifted absorption are highly desired for optogenetic applications. A natural CCR variant with peak absorption in the orange-red spectral region has been discovered and used for photostimulation of neuronal firing (12). However, this CCR (named Chrimson) could not be converted to an anion channel by mutagenesis (13), and no natural ACRs with the absorption maximum beyond 540 nm had been found so far, despite extensive screening of homologous proteins from various cryptophyte species (14, 15). Here we report that among the labyrinthulomycete (“Laby”) ACRs), are four (we name “RubyACRs”) that exhibit maximal spectral sensitivity in the orange-red region of the spectrum, dependent on unique residues of their retinal binding pocket. Further, we also observed energy transfer between a fluorescent protein, fused as a tag to the cytoplasmic C-terminus of a RubyACR, and its photoactive site retinal chromophore, an engineered antenna effect that, to the best of our knowledge, has not been reported in any other microbial rhodopsins.

## Results

### Channelrhodopsin sequences

We found channelrhodopsin homologs in the fully or partially sequenced genomes of nine strains of labyrinthulomycetes from the family *Thraustochytriaceae*. Most of the genomes encode two or three paralogs, with the exception of *Schizochytrium aggregatum* ATCC 28209, in the completely sequenced genome of which only one homolog was detected. The four hits in the *Aurantiochytrium* sp. KH105 genome appear to reflect sequencing/assembly artefacts of only two genes. Some sequences from different organisms were completely or nearly identical to each other. The mean length of the encoded polypeptides was ∼660 residues. The opsin seven transmembrane helical (rhodopsin) domain comprised ∼270 residues and was followed by a large cytoplasmic fragment, as in previously known algal channelrhodopsins. In the cytoplasmic fragment of some homologs the signal receiver domain (REC; accession number cl19078) was detected by bioinformatic analysis, although the E-values were very large (0.01).

We also found six and three channelrhodopsin homologs in the fully sequenced genomes of the haptophytes *Phaeocystis antarctica* and *P. globosa*, respectively. The length of the encoded polypeptides ranges from 312 to 1682 amino acid residues. Besides the rhodopsin domain, no other known conserved domains were detected even in the longest sequence. Our search of the whole-genome shotgun contigs at the National Center for Biotechnology Information (NCBI) also returned two sequences from Stramenopiles sp. TOSAG23-3. Their rhodopsin domains, however, clustered with MerMAIDs, whereas Laby and haptophyte (“Hapto”) homologs formed separate branches of the phylogenetic tree of channelrhodopsins (Fig. 1A).

**Figure 1.**
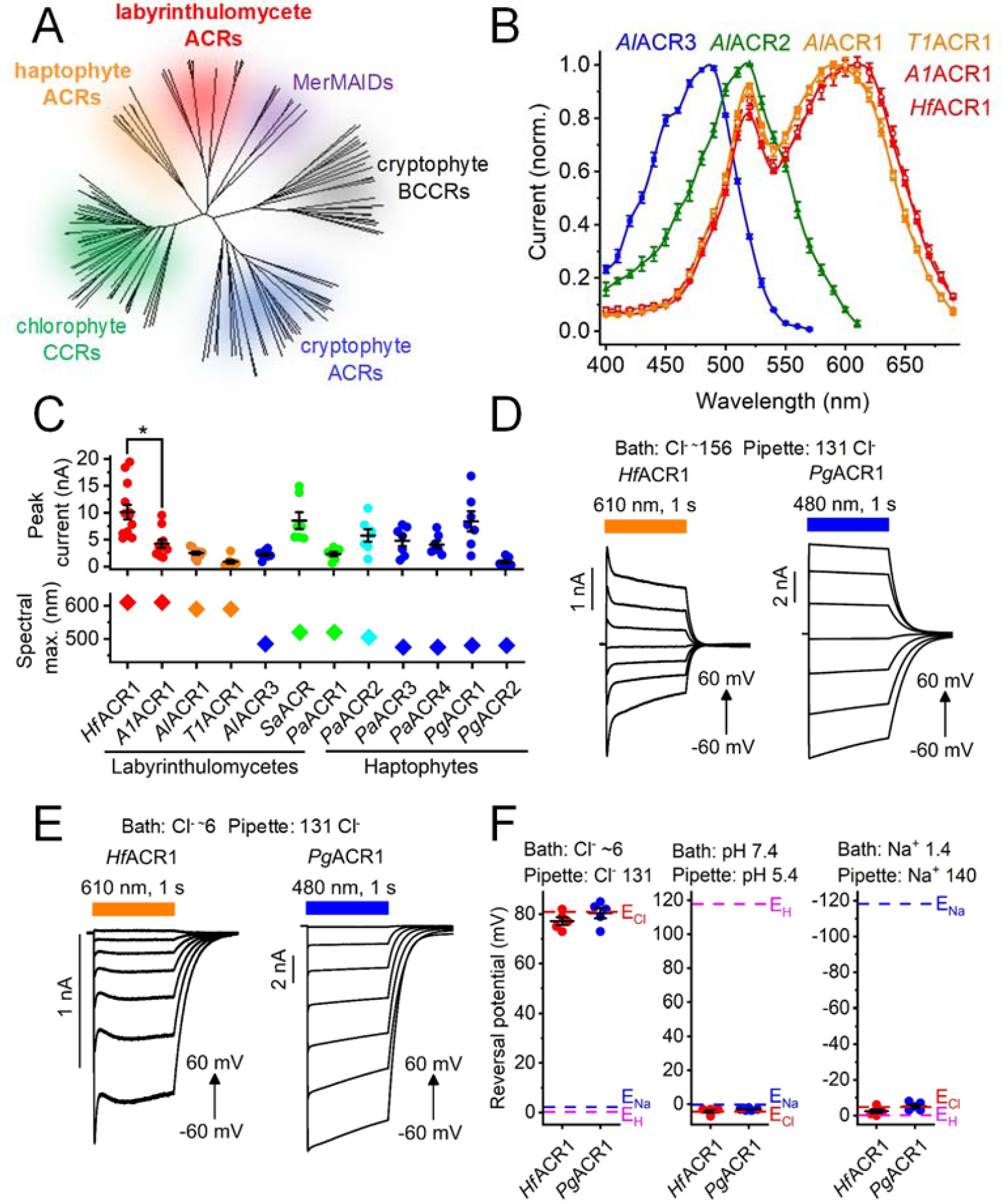
(A) A phylogenetic tree of opsin domains constructed by the neighbor joining method. The GenBank accession numbers and source organism names of the sequences used are listed in Table S1. (B) The action spectra of photocurrents generated by indicated ACRs. (C) The peak current amplitudes (top) and spectral maxima (bottom) of all functional ACRs tested in this study except *Al*ACR2, generated in response to the first 1-s light pulse after seal formation. The data from individual cells are shown as circles, the lines show the mean values ± sem (n = 7-13 cells). (D and E) Series of photocurrents recorded using the indicated Cl^-^ concentrations (in mM) at voltages from −60 to 60 mV at the amplifier output. (F) The reversal potentials (E_rev_) of photocurrents measured under indicated ionic conditions (in mM). The data from individual cells are shown as circles, the lines show the mean values ± sem (n = 5 cells). The E_rev_ values were corrected for liquid junction potentials, as described in Methods.

The GenBank accession numbers, abbreviated protein names, source organism names and gene model numbers of the identified homologs are listed in Tables S1 and S2. The first two italicized letters of the protein names were derived from the genus and species name of the source organism; if no species name was assigned to a strain, consecutive numbers were used to distinguish homologs derived from different strains. When naming homologs from the same organism, we followed a historical numbering convention and assigned the number one to the most red-shifted channelrhodopsin paralog, and consecutively numbered other paralogs according to the position of their spectral maxima (Table S1).

Protein alignments of the rhodopsin domains of Laby and Hapto channelrhodopsins are shown in Figs. S1 and S2, respectively. The Schiff base Lys is found in the seventh transmembrane helix (TM7) of all homologs, as is the immediately following Gln, characteristic of most earlier known channelrhodopsins except BCCRs. The position of the proton donor in bacteriorhodopsin (Asp85) is occupied with a non-carboxylate residue in all the sequences, as in cryptophyte and MerMAIDs ACRs. This feature distinguishes them from chlorophyte CCRs and cryptophyte “bacteriorhodopsin-like” CCRs (BCCRs), most of which contain a carboxylate residue in this position. The second photoactive site carboxylate (Asp212 of bacteriorhodopsin) is universally conserved. Only two glutamates (Glu60 and Glu68 using *Guillardia theta* ACR1 (*Gt*ACR1) numbering) are conserved in TM2 of the Hapto homologs. This is again a feature shared with cryptophyte and MerMAID ACRs, in contrast to chlorophyte CCRs, most of which contain four conserved glutamates in this helix. The labyrinthulomycete homologs completely lack glutamates in TM2.

### Screening of ACR homologs by patch clamp electrophysiology

We synthesized mammalian codon-adapted polynucleotides encoding the rhodopsin domains (residues 1-300) of seven channelrhodopsin homologs from various labyrinthulomycete species, six from *Phaeocystis antarctica* and three from *P. globosa*, fused them to a C-terminal enhanced fluorescent yellow protein (EYFP) tag and expressed them in human embryonic kidney (HEK293) cells. As shown below, Laby and Hapto homologs are strictly anion-selective, so we will refer to them as “ACRs”. Expression of the constructs for *A1*ACR1 from *Aurantiochytrium* sp. and *T1*ACR from *Thraustochytrium* sp. was poor, as judged by the tag fluorescence, but a C-terminal truncation to 270 encoded residues improved it. We also synthesized constructs encoding the entire predicted polypeptides for *Al*ACR1 and *Al*ACR2 from *Aurantiochytrium limacinum* (696 and 635 residues, respectively), but in both cases no membrane fluorescence was observed. The expression construct sequences were deposited to GenBank (their accession numbers are listed in Table S1).

All seven Laby and six Hapto homologs generated photocurrents when probed with whole-cell patch clamping. First, we determined their action spectra by measuring the initial slope in the linear range of the stimulus intensity. The spectra of the three *A. limacinum* homologs peaked at 485, 520 and 590 nm (Fig. 1B), reminiscent of the blue, green and red color system of human vision. *A1*ACR1, *Hf*ACR1 and *T1*ACR1 sequences, although they originate from different organisms, are 76-80% identical to that of *Al*ACR1 (Fig. S1). In the action spectrum of *T1*ACR1 photocurrents the position of the main peak was identical to that of *Al*ACR1 (590 nm), but in both *A1*ACR1 and *Hf*ACR1 it was at 610 nm (Fig. 1B). Other tested homologs exhibited maximal sensitivity to green or blue light. In accordance with a historical numbering convention, we assigned the number one to the most red-shifted channelrhodopsin paralog in a given organism, and consecutively numbered other paralogs according to the position of their spectral maxima (Table S1).

When probed at the wavelength of their maximal sensitivity, most of the tested rhodopsins generated photocurrents in the nA range. Figure 1C shows the peak current amplitude for all tested homologs except *Al*ACR2, the currents of which are described in a separate section below. *Hf*ACR1 generated the largest currents among Laby homologs, and *Pg*ACR1, among the Hapto homologs. Of the two most red-shifted homologs, *Hf*ACR1 generated significantly larger photocurrents than *A1*ACR1 (p < 0.001, Mann-Whitney test), although the sequences of their rhodopsin domains differ only at five positions (Fig. S1). Fig. S3 shows photocurrent traces recorded in response to a 1-s light pulse at −60 mV at the amplifier output from all functional homologs except *A1*ACR2. The amplitude of channel currents decreased under continuous illumination (a phenomenon known as desensitization). The degree of desensitization after 1-s illumination and the half-time of current decay after switching off the light varied between different homologs (Figs. S4A and B). On average, desensitization was greater in Laby homologs (89 ± 3%, mean ± sem, n = 6 homologs) than in Hapto homologs (41 ± 8%, mean ± sem, n = 6 homologs). As in earlier studied strongly desensitizing channelrhodopsins (3), the peak-to-stationary ratio in Laby ACRs reduced in a series of light pulses, whereas in Hapto ACRs this effect was negligible (Fig. S4C).

With standard bath and pipette solutions (∼156 and 131 mM Cl^-^, respectively; for other components see Methods) the photocurrents reversed their direction near zero voltage, indicating passive ionic conductance (Fig. 1D). To test relative permeability for Cl^-^, Na^+^ and H^+^, we individually reduced the concentrations of these ions in the bath. Representative photocurrent traces recorded from *Hf*ACR1 and *Pg*ACR1 with ∼6 mM Cl^-^ in the bath at incremental voltages are shown in Fig. 1E. We then measured the current-voltage relationships and calculated the reversal potentials (E_rev_). Under all tested conditions the E_rev_ followed the Nernst equilibrium potential for Cl^-^ (Fig. 1F), indicating that the tested channelrhodopsins were strictly permeable for anions.

### Color tuning in Laby ACRs

The action spectra of photocurrents generated by all four strongly red-shifted variants (RubyACRs) exhibited an additional sharp peak at 520 nm, suggesting the presence of a second chromophore (Fig. 1A). Comparison of the retinal-binding pockets of *Al*ACR1 (spectral maximum 590 nm) and *Al*ACR3 (487 nm) has revealed four divergent positions (Fig. 2A). Most unusual were Gln213 and Ile217, which correspond, respectively, to Trp182 and Pro186 of bacteriorhodopsin and are highly conserved in the superfamily of microbial rhodopsins. These four unusual residues are conserved in all RubyACRs, but not in Chrimson (Fig. 2A), although the latter also exhibits a red-shifted absorption peak at 590 nm (12).

**Figure 2.**
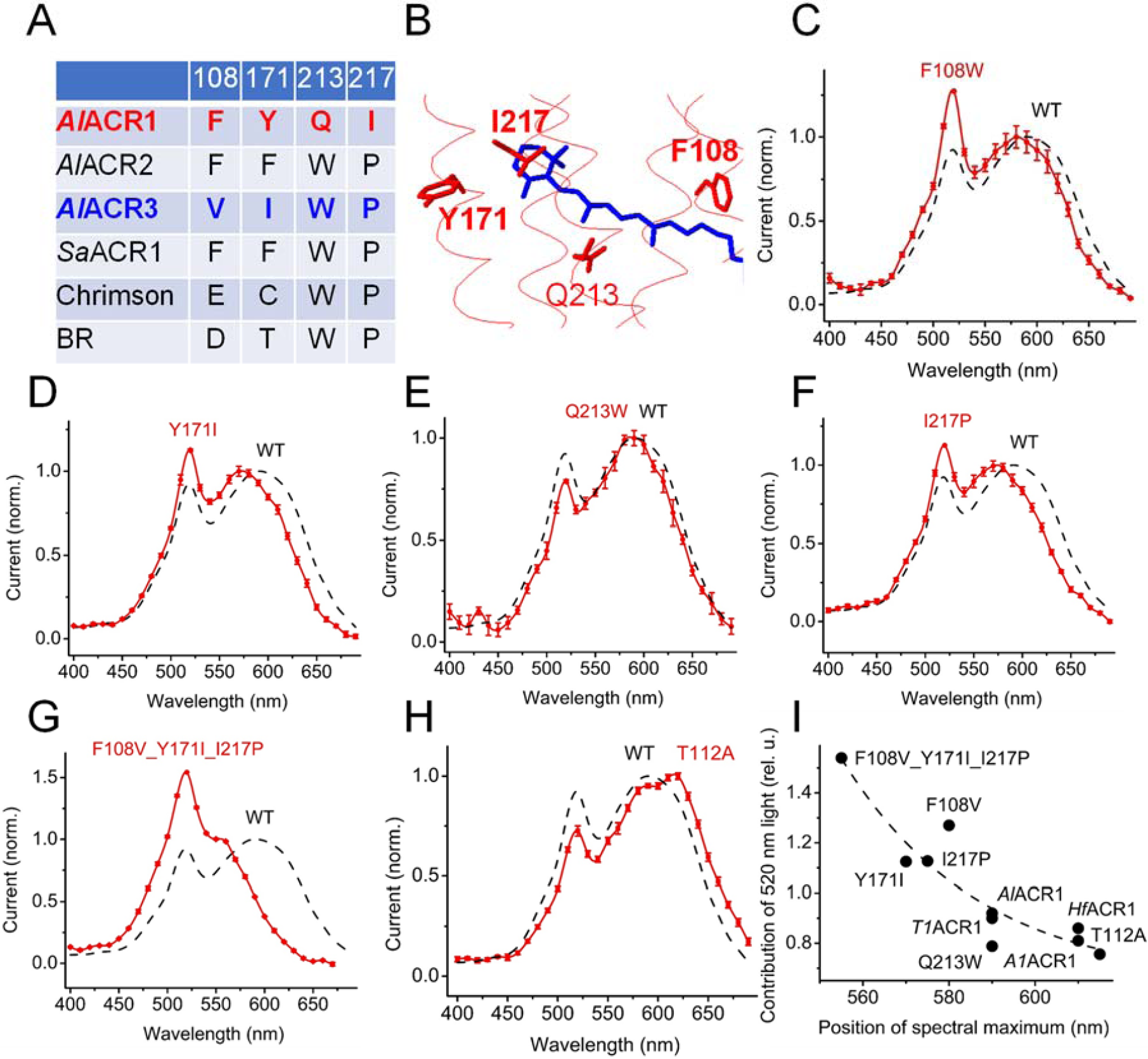
(A) The residues in the retinal binding pocket of indicated proteins. The numbers correspond to the sequence of *Al*ACR1. BR, bacteriorhodopsin. (B) A homology model of *Al*ACR1 showing the side chains of the residues from panel A. (C-H) The action spectra of photocurrents generated by the indicated mutants of *Al*ACR1 (red) and that of the wild type from Fig. 1B (black). (H) The dependence of contribution of the 520-nm peak (its ratio to the rhodopsin peak) on the position of the rhodopsin peak.

Figure 2B shows locations of these residues according to a homology model of *Al*ACR1. To test their possible role in color tuning, we individually replaced the residues in *Al*ACR1 with those found in the corresponding positions in *Al*ACR3 and measured the action spectra of photocurrents in the resultant mutants. In the *Al*ACR1_F108W, Y171I and Y217P mutants the position of the main peak shifted 10, 20 and 15 nm to shorter wavelengths, whereas in the Q213W mutant it remained almost unchanged (Fig. 2C-F). The effect of the three color-tuning mutations was additive (Fig. 2G). In Chrimson the S169A mutation caused a 18-nm shift of the spectral maximum to longer wavelengths (16). The corresponding T112A mutation caused a similar red shift in *Al*ACR1 (Fig. 2H) despite the differences in the retinal binding pockets. In addition to a shift of the main (rhodopsin) spectral peak, the F108W, T112A, Y171I and Y217P mutations also changed the relative contribution of the 520- nm peak (its ratio to the rhodopsin peak). Notably, there was an inverse correlation between this contribution and the rhodopsin peak position (Fig. 2I) as expected from energy migration from the second chromophore to retinal, because efficiency of resonance energy transfer is proportional to the extent of spectral overlap between the donor and acceptor.

Incorporation of A2 retinal (3,4-dehydroretinal) caused a red shift of the spectral sensitivity of several microbial rhodopsins expressed in HEK293 cells (17). Supplementation of *Al*ACR1-expressing cells with A2 retinal led to a red shift of the rhodopsin peak of the action spectrum of photocurrents as compared to that measured with regular (A1) retinal, but did not shift the 520-nm peak (Fig. S5A). In *Hf*ACR1 (the spectral maximum with A1 retinal at 610 nm), not only a red shift of the rhodopsin peak was observed, but also the relative amplitude of the 520-nm peak decreased without a change in its position (Fig. S5B). Both of these effects were clearly resolved in the difference (retinal A2 – retinal A1) spectra (Fig. S5C). These observations confirmed that the 520-nm peak originated from a second chromophore, as its spectral position was not affected by the type of retinal used. Furthermore, energy migration from the antenna pigment to retinal was further confirmed by the inverse correlation of contribution of the antenna peak to the spectrum with the wavelength of the rhodopsin peak, as in the case of the color-tuning mutants (Fig. 2I).

### Retinal-EYFP interaction

To further test whether the second chromophore responsible for the 520-nm peak in the photocurrent action spectra is derived from the EYFP tag and gain more information about the spectral properties of RubyACRs, we expressed *A1*ACR1 and *Hf*ACR1 without fluorescent tags in *Pichia* and purified the encoded proteins in non-denaturing detergent. The absorption spectra of detergent-purified *A1*ACR1 (Fig. 3A) and *Hf*ACR1 (Fig. S6) lacked the 520-nm band observed in the action spectra of the corresponding EYFP fusions. When we replaced the EYFP tag (the absorption maximum at 513 nm) with mCherry tag (587 nm), the 520-nm band disappeared from the action spectrum of photocurrents (Fig. 3A, red). The difference between the action spectrum measured with *Al*ACR1_EYFP and that with *Al*ACR1_mCherry matched the fluorescence excitation spectrum of EYFP (Fig. 3B). The difference between the action spectra also clearly showed that the position of the rhodopsin peak was ∼15-nm shifted to shorter wavelengths by the tag replacement (the arrow in Fig. 3B). The maximum of the action spectrum measured with *Al*ACR1_mCherry was close to that of the absorption spectrum of the fluorescence tag-free protein (595 nm; Fig. 3A black and red).

**Figure 3.**
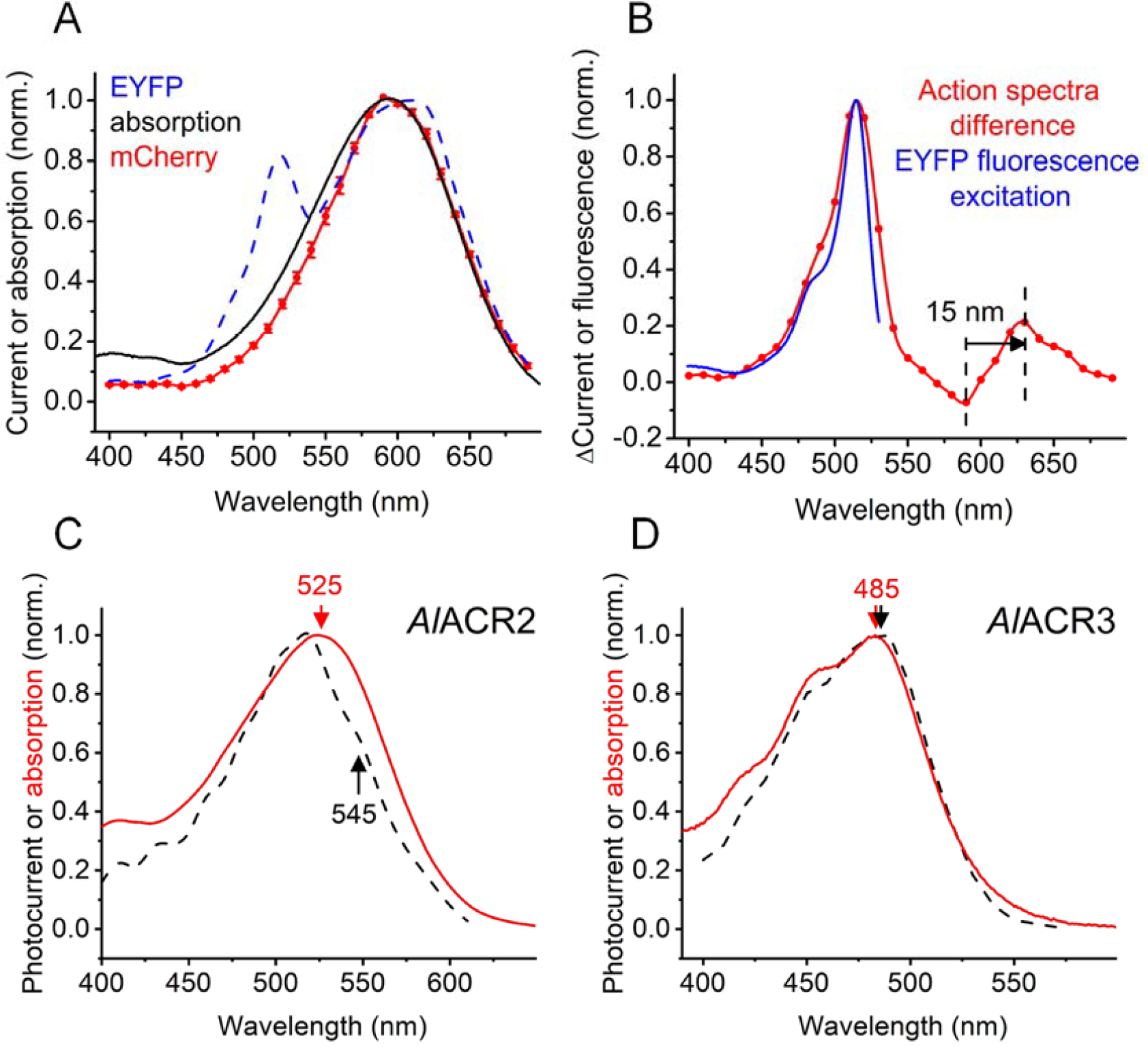
(A) The action spectra of photocurrents generated by *Al*ACR1 fused to EYFP or mCherry and the absorption spectrum of fluorescence tag-free *Al*ACR1 purified from *Pichia*. (B) The difference between the action spectra obtained with *Al*ACR1_EYFP and *Al*ACR1_mCherry and the fluorescence excitation spectrum of EYFP from FPbase (https://www.fpbase.org/protein/eyfp/). (C and D) The action spectra of photocurrents recorded by *Al*ACR2 (C) and *Al*ACR3 (D) EYFP fusions (black dashed lines, reproduced from Fig. 1B) and the absorption spectra of the respective purified proteins (solid red lines).

The absorption spectrum of the *Pichia*-expressed green-absorbing *Al*ACR2 peaked at 525 nm (Fig. 3C, red). The action spectrum of photocurrents recorded from the *Al*ACR2_EYFP fusion exhibited a shoulder at ∼545 nm (Fig. 3C, black). Comparison with the absorption spectrum show that this shoulder corresponds to rhodopsin absorption, whereas the main peak at 520 nm reflects energy transfer from EYFP. The absorption spectrum of the *Pichia*-expressed blue-absorbing *Al*ACR3 closely matched the action spectrum of photocurrents recorded from the corresponding EYFP fusion (Fig. 3D).

### Channel gating in AlACR1

To analyze the kinetics of channel gating in *Al*ACR1, we recorded photocurrents under single-turnover conditions using 6-ns laser flash excitation. Laser-evoked current traces could be fit with four exponentials (two for the current rise and two for the decay) (Fig. 4A). Channel opening (the time constant (τ) ∼100 µs) was preceded with a fast, only weakly voltage-dependent negative peak that could be clearly resolved near the reversal voltage for channel currents (the arrow in Fig. 4A). Its rise τ was <20 µs, which was the lower limit of the time resolution of our system. Such currents have previously been recorded from several other channelrhodopsins and attributed to a charge displacement associated with retinal isomerization, integrated by the recording circuit (18).

**Figure 4.**
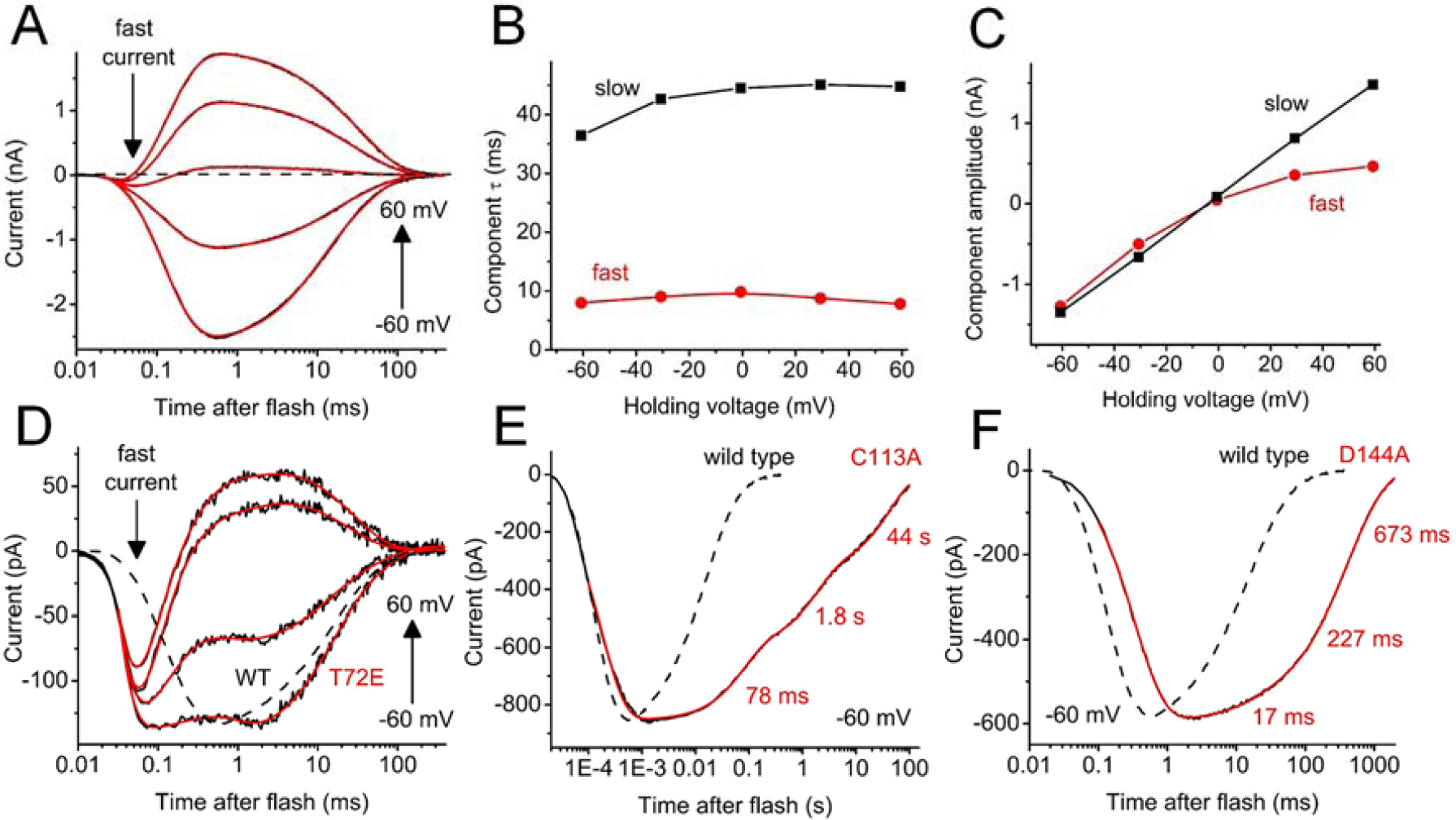
(A) A series of photocurrent traces recorded from *Al*ACR1 upon laser flash excitation at the voltages changed in 20 mV increment from −60 to 60 mV at the amplifier output. Black lines, experimental data; red lines, multiexponential fit. (B and C) The dependence of the decay components τ (B) and amplitude (C) on the holding voltage for the series of traces shown in panel A. (D) A series of photocurrent traces recorded from the *Al*ACR1_T72E mutant at incremental voltages from −60 to 60 mV. The wild-type (WT) trace at −60 mV from panel A is shown as the dashed line for comparison. (E and F) The current traces recorded from the indicated *Al*ACR1 mutants at −60 mV (solid lines) as compared to the wild-type trace (dashed lines).

*Al*ACR1 channel closing was biphasic, and both phases were faster (τ 13 ± 1 ms and 77 ± 11 ms at −60 mV, n = 5 cells) than those in *Gt*ACR1 (19). Moreover, in *Al*ACR1 τ of both decay phases showed only weak (if any) voltage dependence (Fig. 4B), in contrast to *Gt*ACR1 in which the fast decay was strongly accelerated, and the slow decay slowed upon depolarization (19). The amplitude of the fast decay component in *Al*ACR1 exhibited inward, and that of the slow decay, no rectification (Fig. 4C), again in contrast to *Gt*ACR1, in which the amplitude of the fast component showed outward, and that of the slow component, inward rectification (19).

Previously we had found that Glu68 controls the kinetics of the fast channel closing in *Gt*ACR1 (19). The corresponding position in *Al*ACR1 is occupied by a non-carboxylate residue, Thr72 (Fig. S1), which is very unusual for channelrhodopsins, in most of which this residue is conserved. The T72E mutation significantly reduced laser-evoked channel currents from 2.7 ± 0.5 nA measured in the wild type to 76 ± 17 pA (mean ± s.e.m., n = 8 and 7 cells, respectively; p < 0.005, Mann-Whitney test), which made the fast isomerization peak more obvious in the current trace (Fig. 4D). The current decay rate was little affected by this mutation. Cys128 from TM3 and Asp156 from TM4 form an interhelical hydrogen bond (the “DC gate”) in *Chlamydomonas reinhardtii* channelrhodopsin-2 (*Cr*ChR2) (20). The homologs of these residues in *Al*ACR1 are Cys113 and Asp144. Alanine replacement of each of these residues slowed channel closing (Fig. 4E and F). The effect was much more pronounced in the C113A mutant (the slowest decay phase was on the time scale of tens of seconds) than in the D144A mutant.

### Unusual photocurrents of AlACR2

In contrast to all other channelrhodopsins studied in heterologous expression systems amenable to electrophysiological recording, photocurrents from *Al*ACR2 were dominated by a fast negative signal (Fig. 5A). Its peak amplitude recorded in response to a 1-s pulse of continuous light at −60 mV was 103 ± 15 pA (mean ± sem, n = 7 cells). It showed a very weak, in any, dependence of the holding voltage (Fig. 5B), as is typical of similar fast negative currents associated with retinal isomerization in other channelrhodopsins (18). To detect a possible contribution of passive Cl^-^ conductance in photocurrents, we reduced the Cl^-^ concentration in the pipette to 4 mM and held the cells at +60 mV. Under these conditions, if there were any passive Cl^-^ current, it would be outward (i.e., Cl^-^ influx), opposite to the inward isomerization current. The laser flash-evoked current traces could be fit with four exponentials, the slowest of which had τ 10 ms (Fig. 5C), which is within the range of channel closing time in other ACRs. Next, we measured the voltage dependence of the amplitude of this component with 4 and 131 mM Cl^-^ in the pipette. Fig. 5D shows that, in contrast to the peak amplitude, the reversal potential of this slowest component followed the equilibrium potential for Cl^-^, which confirmed that *Al*ACR2 possessed residual passive conductance for this ion.

**Figure 5.**
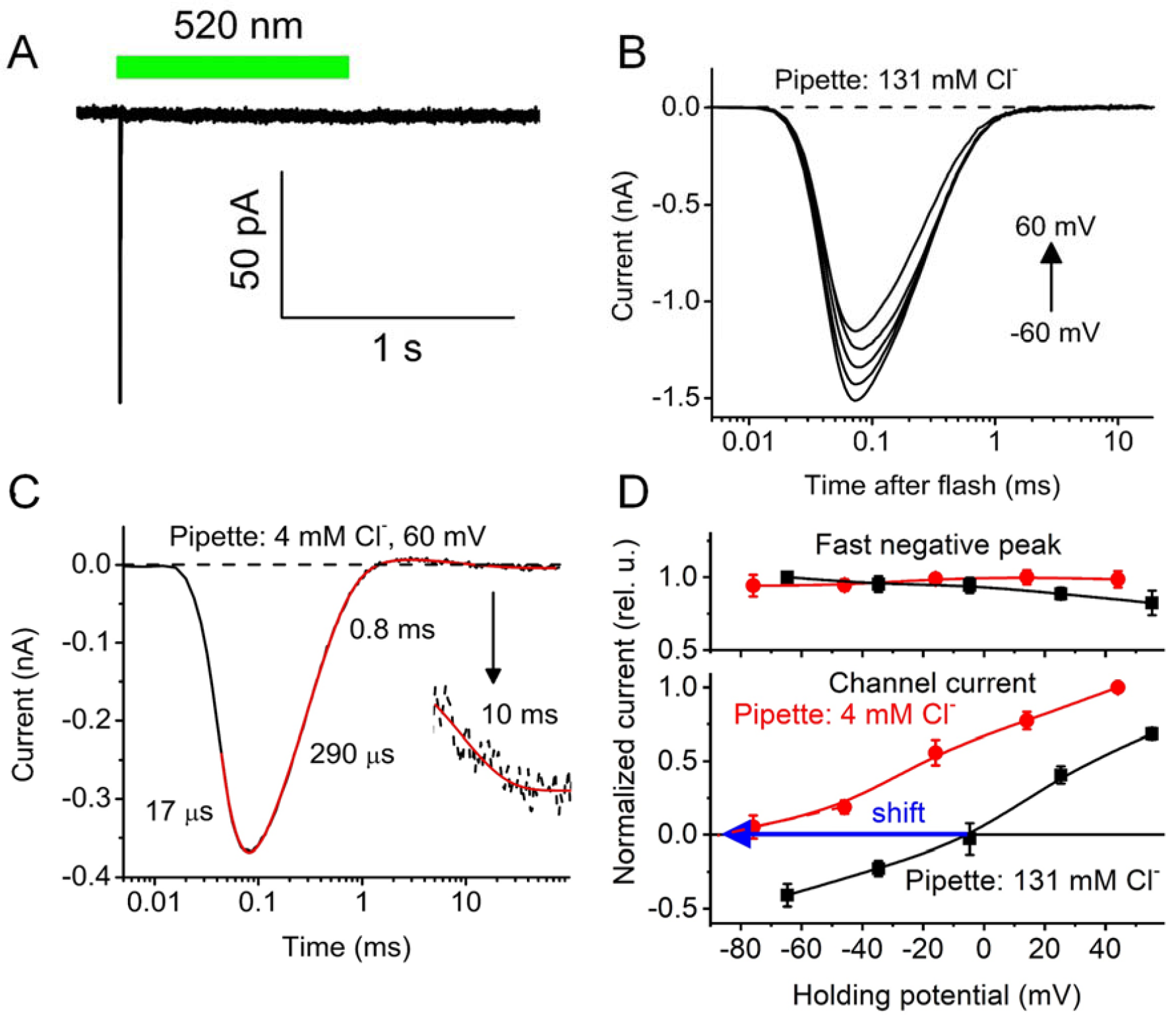
(A) A photocurrent trace recorded from *Al*ACR2 upon stimulation with a continuous light pulse, the duration of which is schematically shown as the green bar. (B) A series of photocurrent traces recorded from *Al*ACR2 upon laser flash excitation at the voltages changed from −60 to 60 mV at the amplifier output with the standard pipette solution. (C) Main figure: A photocurrent trace recorded from *Al*ACR2 at 60 mV using the pipette solution with a reduced Cl^-^ concentration. Inset: a portion of the trace from 5 to 100 ms vertically stretched to better resolve the channel current decay. Black lines, experimental data; red lines, multiexponential fit. (D) The current-voltage dependencies of the peak current and the slowest signal component at the indicated Cl^-^ concentrations in the pipette. The data points are the mean ± sem values (n = 5 cells).

The action spectrum of the unusual photocurrents generated by the *Al*ACR2_EYFP fusion exhibited a main peak at 520 nm and a minor shoulder at ∼540 nm (Fig. 1B, green). The position of the main peak was very close to that of EYFP absorption. To test the possibility of energy transfer from EYFP to the *Al*ACR2 retinal chromophore, we measured the action spectrum after incubation of the cells with A2 retinal. A clear shoulder appeared at ∼580 nm, which we interpret as likely due to a shift of the rhodopsin peak to longer wavelengths, whereas the peak at 520 nm remained unchanged (Fig. S5D) consistent with energy transfer between EYFP and *Al*ACR2.

## Discussion

We report channelrhodopsins in two eukaryotic lineages phylogenetically distant from green and cryptophyte algae in which such proteins have been found so far. Both Laby and Hapto ACRs conduct exclusively anions and therefore should be classified as ACRs. This expansion of the ACR family benefits elucidation of the structural requirements for anion conductance by comparative analysis of the structure-function relationships in different ACR groups and offers a possibility of using alternative scaffolds for molecular design of physico-chemical properties desirable in optogenetic tools. The most important in this respect are the four RubyACRs, closely related Laby ACRs with spectral maxima in the orange-red region. The only so far known inhibitory optogenetic tool with the peak absorption at ∼590 nm has been the engineered halorhodopsin from *H. salinarum* strain Shark, referred to as Jaws (21). Although Jaws generates larger photocurrents than the earlier known Cl^-^ pumps, its efficiency is limited by translocation of only one ion per captured photon across the membrane. RubyACRs generate large passive anion currents and offer a new type of highly efficient optogenetic inhibition for the long-wavelength spectral range. Our analysis of spectral-tuning mutants and A1/A2 retinal substitution strongly indicate Förster resonance energy transfer from a fluorescent protein (EYFP) and retinal in red-shifted rhodopsins, which may provide an additional avenue for the development of optogenetic tools.

A conserved amino acid residue pattern of the retinal binding pocket in RubyACRs has not been found in any other microbial rhodopsins, including the cation-conducting Chrimson that exhibits peak absorption at 590 nm. Of the three pocket residues that contribute to the unique spectral sensitivity of RubyACRs, Phe108 (*Al*ACR1 numbering) occupies the position of the counterion of the Schiff base (Asp85 in bacteriorhodopsin), the importance of which for color tuning is well documented (22, 23). As shown in bacteriorhodopsin, a neutral residue in this position leads to a decrease in the energy gap between the ground and excited states resulting in red-shifted absorption (24). A bulky aromatic residue in the counterion position is very unusual among microbial rhodopsins. A previously known group of sequences in which phenylalanine was found are schizorhodopsins from Asgardarchaeota, but they show very little homology to channelrhodopsins (25). To the best of our knowledge, a possible role of this substitution in color tuning in schizorhodopsins has not been tested. According to our homology model of *Al*ACR1, Tyr171 and Ile217 are located near the β-ionone ring. The effect of the polar Tyr171 on the *Al*ACR1 spectrum is likely due to stabilization of the excited state of the chromophore, as has been shown for the more polar ring environment in bacteriorhodopsin, as compared to that in sensory rhodopsin II (26).

The only species in which the functional role of channelrhodopsins has been demonstrated is the green flagellate alga *Chlamydomonas reinhardtii*, in which they function as phototaxis receptors (1). All labyrintholomycete and haptophyte species in which we found channelrhodopsins produce free-swimming zoospores in their lifecycles (27, 28). Therefore, it is plausible that channelrhodopsins guide phototaxis also in these microorganisms. Consistent with this hypothesis is that no channelrhodopsin homologs have been found in the fully sequenced genome of the labyrintholomycete *Aplanochytrium kerguelense* PBS07, which does not produce zoospores (28).

## Materials and Methods

### Bioinformatics

A keyword search of gene annotations was used to identify rhodopsin genes at the genome portals of *A. limacinum* MYA-1381, *P. antarctica* CCMP1374 and *P. globosa* Pg-G of the US Department of Energy JGI. The three hits showing protein sequence homology to previously known algal channelrhodopsins found in the *A. limacinum* genome were then used for tblastn search of the JGI genome portal for *Schizochytrium aggregatum* ATCC 28209 and whole-genome shotgun contigs from stramenopiles at the NCBI portal.

Protein sequence alignments were created using MUSCLE algorithm implemented in DNASTAR Lasergene (Madison, WI) MegAlign Pro software. Phylogenetic trees were visualized using Dendroscope software (29). A homology model of *Al*ACR1 was obtained using i-TASSER server (30) and visualized by PyMol software (http://www.pymol.org).

### Molecular biology

For expression in HEK293 cells, DNA polynucleotides encoding the transmembrane domains showing homology to previously known ACRs optimized for human codon usage were synthesized (GenScript, Piscataway, NJ) and cloned into the mammalian expression vector pcDNA3.1 (Life Technologies, Grand Island, NY) in frame with an EYFP or mCherry tag. Mutants were generated using Quikchange XL kit (Agilent Technologies, Santa Clara, CA) and verified by sequencing. For expression in *Pichia*, the opsin-encoding constructs were fused in frame with a C-terminal 8-His tag and subcloned into the pPIC9K (*A1*ACR1, *Hf*ACR1 and *Al*ACR2) or pPICZα (*Al*ACR3) vector (Invitrogen) according to the manufacturer’s instructions.

### HEK293 transfection and patch clamp recording

HEK293 cells were transfected using the JetPRIME transfection reagent (Polyplus, Illkirch, France). All-*trans*-retinal (Sigma) was added at the final concentration of 3 µM immediately after transfection. Photocurrents were recorded 48-96 h after transfection in the whole-cell voltage clamp mode with an Axopatch 200B amplifier (Molecular Devices, Union City, CA) using the 10 kHz low-pass Bessel filter. The signals were digitized with a Digidata 1440A using pClamp 10 software (both from Molecular Devices). Patch pipettes with resistances of 2-4 MΩ were fabricated from borosilicate glass. The standard pipette solution contained (in mM): KCl 126, MgCl_2_ 2, CaCl_2_ 0.5, Na-EGTA 5, HEPES 25, pH 7.4. The standard bath solution contained (in mM): NaCl 150, CaCl_2_ 1.8, MgCl_2_ 1, glucose 5, HEPES 10, pH 7.4. A 4 M KCl bridge was used in all experiments, and possible diffusion of Cl^-^ from the bridge to the bath was minimized by frequent replacement of the bath solution with fresh buffer. For measurements of the reversal potential shifts under varied ionic conditions, Na^+^ was substituted for K^+^ in the pipette solution to minimize the number of ionic species in the system. To reduce the Cl^-^ concentration in the bath, NaCl was replaced with Na-aspartate; to reduce the Na^+^ concentration, with N-methyl-D-glucamine chloride; to increase the H^+^ concentration, pH was adjusted with H_2_SO_4_. The holding voltages were corrected for liquid junction potentials calculated using the Clampex built-in LJP calculator (31). Continuous light pulses were provided by a Polychrome V light source (T.I.L.L. Photonics GMBH, Grafelfing, Germany) in combination with a mechanical shutter (Uniblitz Model LS6, Vincent Associates, Rochester, NY; half-opening time 0.5 ms). The action spectra were constructed by calculation of the initial slope of photocurrent and corrected for the quantum density measured at each wavelength. Laser excitation was provided by a Minilite Nd:YAG laser (532 nm, pulsewidth 6 ns, energy 12 mJ; Continuum, San Jose, CA). The current traces were logarithmically filtered using a custom software. Curve fitting was performed by Origin Pro software (OriginLab Corporation, Northampton, MA).

### Expression and purification of ACRs from Pichia

The plasmids encoding Laby ACRs were linearized with SalI or PmeI and used to transform *P. pastoris* strain SMD1168 (*his4, pep4*) by electroporation according to the manufacturer’s instructions. Resistant transformants were screened on 4 mg/ml geneticin or 1 mg/ml zeocin and first cultivated on a small scale. Rhodopsin gene expression was induced by the addition of methanol. All-*trans* retinal (5 µM final concentration) was added simultaneously. Clones of the brightest color were selected for further experimentation. For protein purification, a starter culture was inoculated into buffered complex glycerol medium until A600 reached 4–8, after which the cells were harvested by centrifugation and transferred to buffered complex methanol medium supplemented with 5 μM all-*trans* retinal (Sigma Aldrich). Expression was induced by the addition of 0.5% methanol. After 24-30 h, the cells were harvested and disrupted in a bead beater (BioSpec Products, Bartlesville, OK) in buffer A (20 mM sodium phosphate, pH 7.4, 100 mM NaCl, 1 mM EDTA, 5% glycerol). After removing cell debris by low-speed centrifugation, membrane fragments were collected by ultracentrifugation, resuspended in buffer B (20 mM Hepes, pH 7.4, 300 mM NaCl, 5% glycerol) and solubilized by incubation with 1.5% dodecyl maltoside (DDM) for 1.5 h or overnight at 4°C. Non-solubilized material was removed by ultracentrifugation, and the supernatant was mixed with nickel-nitrilotriacetic acid or cobalt superflow agarose beads (Thermofisher) and loaded on a column. The proteins were eluted with buffer C (20 mM Hepes, pH 7.4, 300 mM NaCl, 5% glycerol, 0.02% DDM) containing 300 mM imidazole, which was removed by repetitive washing with imidazole-free buffer C using YM-10 centrifugal filters (Amicon, Billerica, MA).

### Absorption spectroscopy

Absorption spectra of purified proteins were recorded using a Cary 4000 spectrophotometer (Varian, Palo Alto, CA).

### Statistics

The data are presented as mean ± s.e.m. values. Normality and equal variances of the data were not assumed, and therefore the non-parametric two-sided Mann-Whitney test was used for pairwise comparison of independent data sets. P values > 0.05 were considered not significant. The sample size was estimated from previous experience and published work on a similar subject, as recommended by the NIH guidelines (32).

## Acknowledgments

This work was supported by the National Institutes of Health Grant R01GM027750, the Hermann Eye Fund, and Endowed Chair AU-0009 from the Robert A. Welch Foundation to J.L.S, and by the Natural Sciences and Engineering Research Council of Canada (NSERC) Discovery Grant RGPIN-2018-04397 to L.S.B.

## Notes

### Competing Interest Statement

The authors have declared no competing interest.

