## Supplementary Material for "RubyACRs, non-algal anion channelrhodopsins with highly red-shifted absorption"

* Address correspondence to: John L. Spudich.

**This PDF file includes:**

Figures S1 to S6
Tables S1 to S2

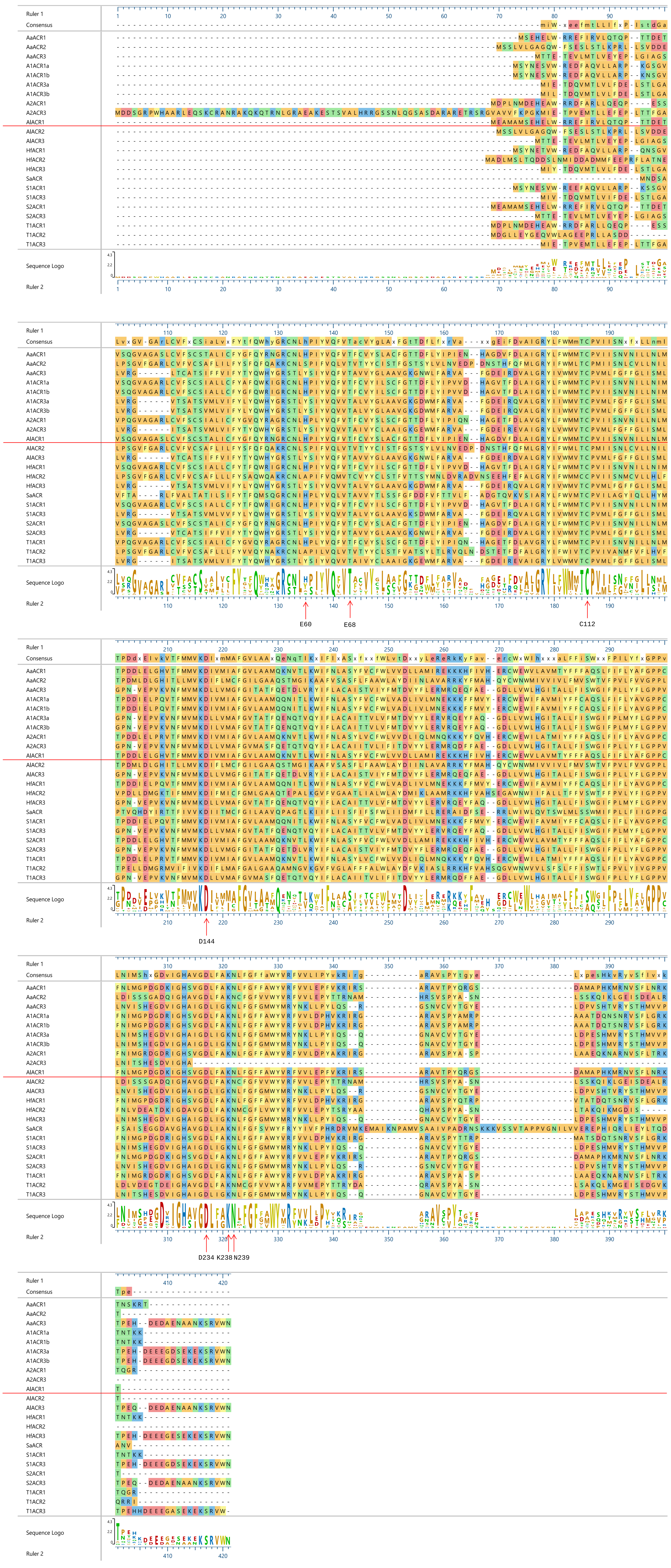
Ruler 1
Consensus
AaACR1
AaACR2
AaACR3
A1ACR1a
A1ACR1b
A1ACR3a
A1ACR3b
A2ACR1
A2ACR3
AIACR1
AIACR2
AIACR3
HfACR1
HfACR2
HfACR3
SaACR
S1ACR1
S1ACR3
S2ACR1
S2ACR3
T1ACR1
T1ACR2
T1ACR3
Sequence Logo
Ruler 2
E60
E68
C112
D144
D234
K238
N239

**Fig. S1.** Protein sequence alignment of the rhodopsin domains of Laby channelrhodopsins. Residues are color-coded according to their chemical properties. The arrows point to the positions of the residues known to be functionally important in *Gt*ACR1.

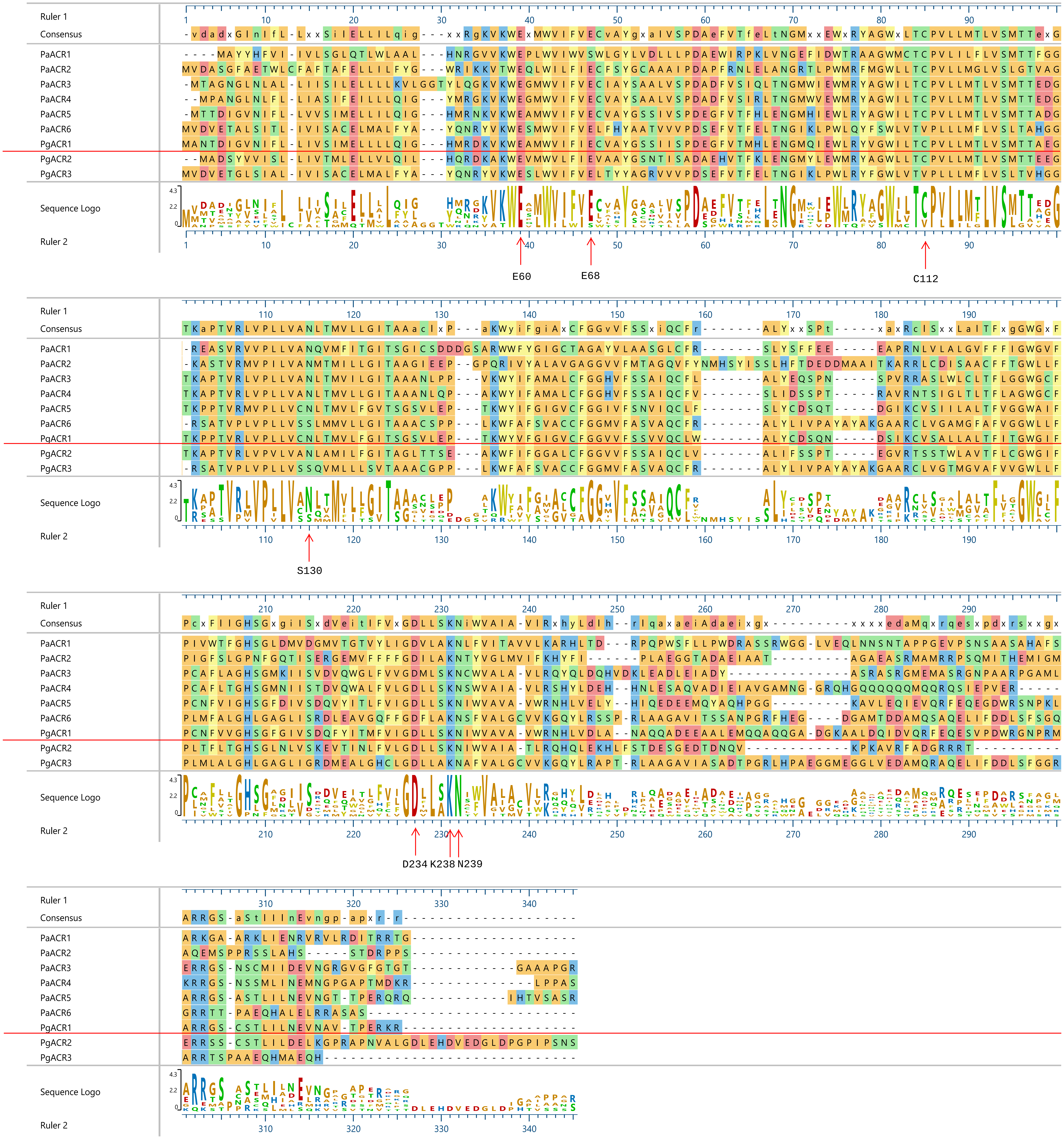
Ruler 1
Consensus
PaACR1
PaACR2
PaACR3
PaACR4
PaACR5
PaACR6
PgACR1
PgACR2
PgACR3
Sequence Logo
Ruler 2
E60
E68
C112
S130
D234
K238
N239

**Fig. S2.** Protein sequence alignment of the rhodopsin domains of Hapto channelrhodopsins. Residues are color-coded according to their chemical properties. The arrows point to the positions of the residues known to be functionally important in *Gt*ACR1.

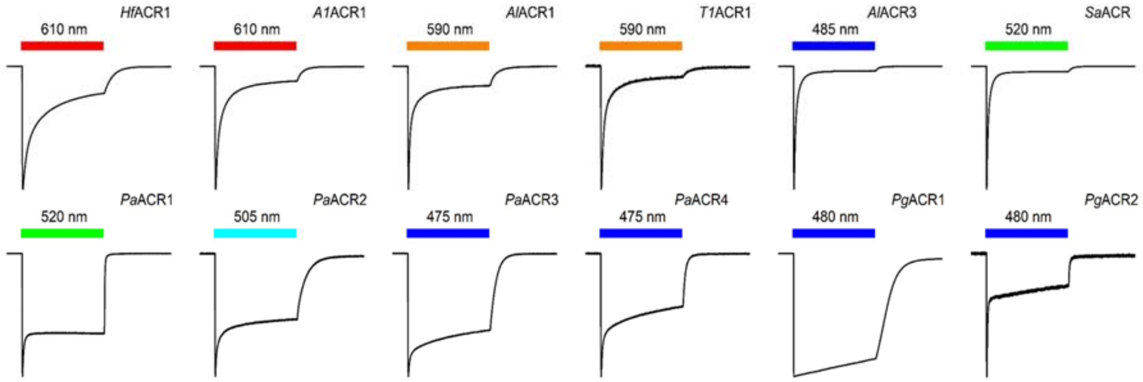


**Fig. S3.** Photocurrent traces recorded in response to the first 1-s light pulse at -60 mV at the amplifier output, normalized at their peak value. The duration of illumination is showed as a colored bar on top.

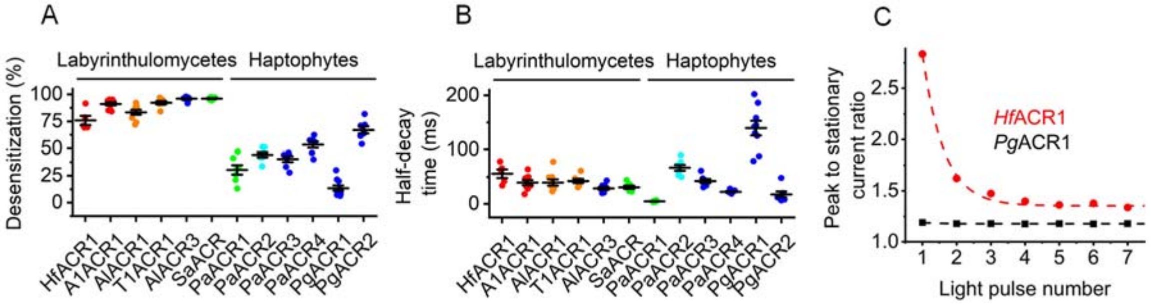


**Fig. S4.** (A and B) The magnitude of desensitization during continuous illumination (A) and half-decay of photocurrent after switching the light off (B). The black lines show the mean values and s.e.m. (n = 5-10 cells for each variant); colored circles, the individual data points. (C) The ratio of the peak amplitude to that of the stationary current (measured at the end of a 1-s light pulse) in a series of pulses applied with 30-s time interval. The lines are single exponential fits.

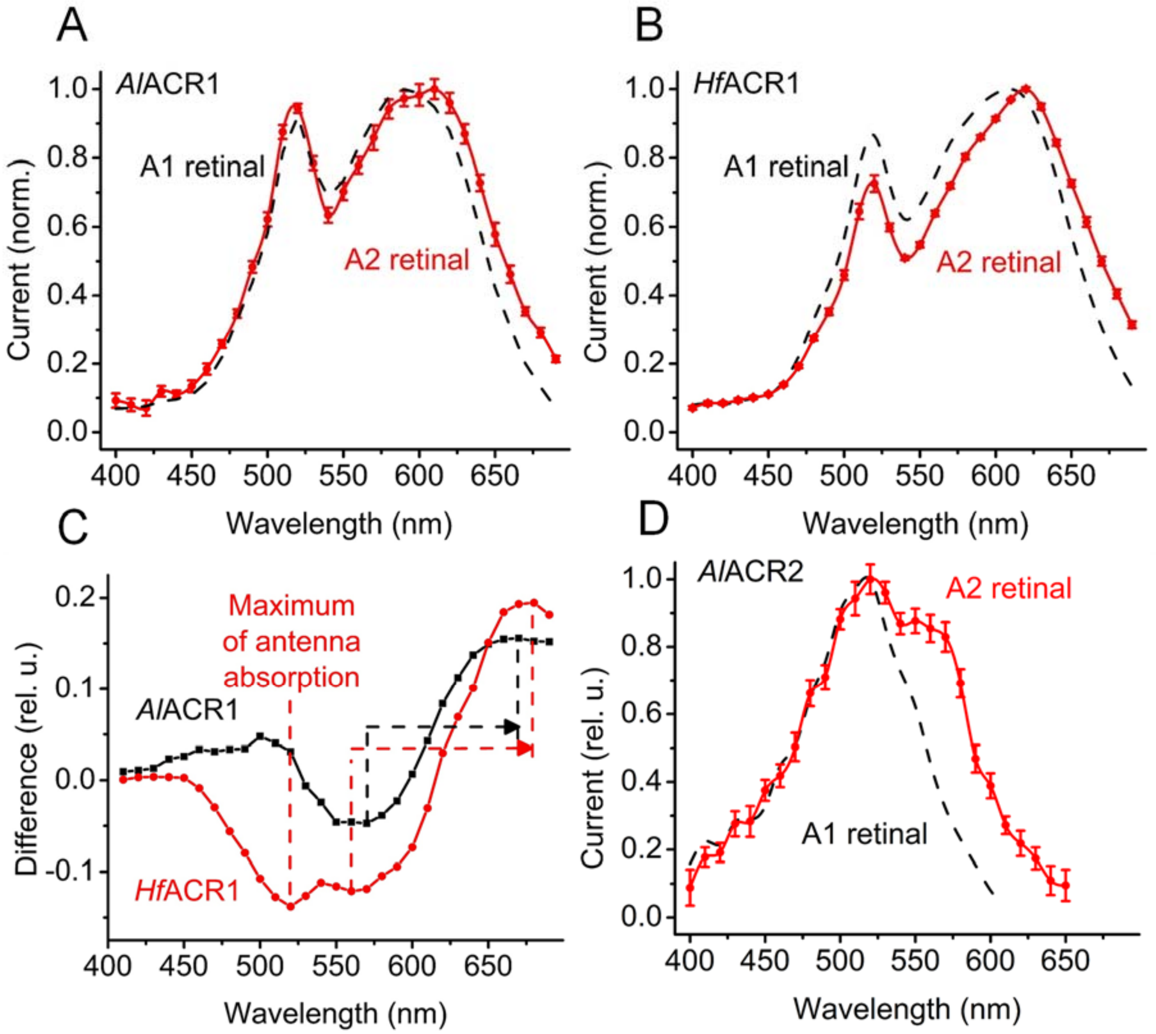


**Fig. S5.** (A, B and D) The action spectra of photocurrents generated by indicated proteins reconstituted with A1 (black) and A2 (red) retinal. (C) The difference spectra (A2-A1 retinal).

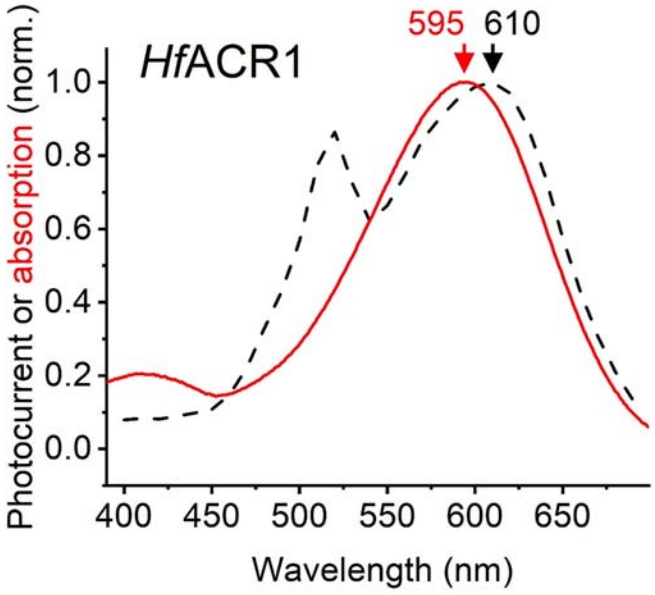


**Fig. S6.** The absorption spectrum of *Hf*ACR1 detergent-purified from *Pichia* (red solid line) compared to the action spectrum of photocurrents generated upon its expression in HEK293 cells from Fig. 1B (black dashed line).

**Table S1.** A list of Laby ACR homologs (in bold – synthesized and tested by patch clamp in this study)

| | GenBank accession number | Abbreviated protein name | Source organism | JGI gene model name | Total CDS length | $\lambda_{max}$ (nm) |
|---|---|---|---|---|---|---|
| 1. | QDJC01000532 | *Aa*ACR1 | *Aurantiochytrium acetophilum* HS399 | identical to *Al*ACR1 | 691 | |
| 2. | QDJC01003161 | *Aa*ACR2 | | identical to *Al*ACR2 | 635 | |
| 3. | QDJC01000037 | *Aa*ACR3 | | only two mismatches with *Al*ACR3 | 683 | |
| **4.** | **MT002467** | ***Al*ACR1** | ***Aurantiochytrium limacinum* ATCC MYA-1381** | **fgenesh1_pg.12_#_284** | **696** | **590** |
| **5.** | **MT002473** | ***Al*ACR2** | | **gm1.7690_g** | **635** | **545** |
| **6.** | **MT002476** | ***Al*ACR3** | | **estExt_Genemark1.C_1_t20010** | **678** | **485** |
| **7.** | **MT002468, BGKB01000037** | ***A1*ACR1** | ***Aurantiochytrium* sp. KH105** | | **646** | **610** |
| **8.** | BGKB01000105 | *A1ACR1* | | | 645 | |
| **9.** | BGKB01000099 | *A1*ACR3 | | | 680 | |
| **10.** | BGKB01000102 | *A1*ACR3 | | | 680 | |
| **11.** | LNGJ01004228 | *A2*ACR1 | *Aurantiochytrium* sp. T66 | identical to *T1*ACR1 | 649 | |
| **12.** | LNGJ01002066 | *A2*ACR3 | | | 759 | |
| **13.** | **MT002469, GBG24568** | ***Hf*ACR1** | ***Hondaea fermentalgiana* FCC1311** | | **646** | **610** |
| **14.** | GBG24569 | *Hf*ACR2 | | | 663 | |
| **15.** | GBG23965 | *Hf*ACR3 | | | 680 | |
| **16.** | **MT002463** | ***Sa*ACR** | ***Schizochytrium aggregatum* ATCC 28209** | **fgenesh1_pg.3_#_476** | **546** | **520** |
| 17. | JTFK01000019 | *S1*ACR1 | *Schizochytrium sp.* CCTCC M209059 | | 645 | |
| 18. | JTFK01000324 | *S1*ACR3 | | | 680 | |
| 19. | SMSO01000032 | *S2*ACR1 | *Schizochytrium sp.* TIO01 | Identical to *Al*ACR1 | 696 | |
| 20. | SMSO01000014 | *S2*ACR3 | | | 583 | |
| **21.** | **MT002470, MUFY01006470** | ***T1*ACR1** | ***Thraustochytrium* sp. ATCC 26185** | | **649** | **590** |
| 22. | MUFY01006469 | *T1*ACR2 | | | 666 | |
| **23.** | MUFY01009420 | *T1*ACR3 | | | 682 | |

**Table S2.** A list of Hapto ACR homologs tested in this study

| | GenBank accession number | Abbreviated protein name | Source organism | JGI gene model name | Total CDS length | $\lambda_{max}$ (nm) |
|---|---|---|---|---|---|---|
| 1. | MT002471 | *Pa*ACR1 | *Phaeocystis antarctica* CCMP1374 | Phant.0066s0015.1 | 1682 | 520 |
| 2. | MT002474 | *Pa*ACR2 | | Phant.0011s0329.1 | 647 | 505 |
| 3. | MT002477 | *Pa*ACR3 | | Phant.0016s0461.1, Phant.0016s0462.1, Phant.0016s0464.1 | 427 | 475 |
| 4. | MT002464 | *Pa*ACR4 | | Phant.0060s0074.1 | 469 | 475 |
| 5. | MT002465 | *Pa*ACR5 | | Phant.0086s0086.1 | 312 | N.A. |
| 6. | MT002466 | *Pa*ACR6 | | Phant.0001s0932.1 | 471 | N.A. |
| 7. | MT002472 | *Pg*ACR1 | *Phaeocystis globosa* Pg-G | Phglo.0395s0005.1 | 327 | 480 |
| 8. | MT002475 | *Pg*ACR2 | | Phglo.0149s0014.1 | 435 | 480 |
| 9. | MT002478 | *Pg*ACR3 | | Phglo.0128s0040.1 | 505 | N.A. |
